# Joint Representation of Valence and Arousal in the Human Brain

**DOI:** 10.64898/2026.11.24.768770

**Authors:** Yujun Chen, Andreas Keil, Mingzhou Ding

## Abstract

Scenes varying in emotional content are commonly characterized along two affective dimensions: hedonic valence, reflecting pleasantness, and emotional arousal, reflecting intensity. Although prior studies have examined neural representations of valence and arousal separately, it remains unclear how these dimensions are represented jointly and how their representation relates to low-level visual and scene-level properties. Here, we used representational similarity analysis (RSA) to identify brain regions whose activity patterns represented the joint valence–arousal structure of affective images and to compare this representation with those of low-level visual and scene-level properties. Twenty participants viewed 60 International Affective Picture System (IAPS) images during simultaneous EEG–fMRI recording. We constructed three image-level representational dissimilarity matrices (RDMs): an Emotion RDM derived from valence and arousal ratings; a GIST RDM derived from low-level visual features captured by GIST descriptors; and a Scene RDM derived from CLIP-ViT image embeddings. Searchlight fMRI RSA identified joint valence–arousal representations in lateral occipitotemporal cortex, low-level visual representations in visual cortex, with smaller extensions into right STS and right insula, and scene-level representations across distributed occipital, temporal, parietal, and frontal regions. The valence–arousal representation showed little spatial overlap with the low-level visual representation but was spatially nested within regions representing scene-level properties. EEG–fMRI fusion further indicated that low-level visual representations emerged at approximately 124 ms after stimulus onset, followed by scene-level and valence–arousal representations at approximately 428 ms and 532 ms, respectively. Together, these findings suggest that joint valence–arousal representations are linked to higher-order scene representations, are distinct from low-level visual representations in both space and time, and may support the affective-semantic characterization of visual scenes. Exploratory analyses further showed that valence and arousal ratings generated by multimodal large language models (ChatGPT, Claude, and Gemini) corresponded closely with human ratings and yielded neural representation maps that substantially overlapped with the human valence–arousal map.

## Introduction

The ability to extract affective information from complex visual scenes is an important function of the human visual system. This process is commonly studied by presenting observers with images of natural scenes varying in affective content. Affective scenes are typically characterized along two dimensions: valence, or pleasantness, and arousal, or intensity. A central question in affective neuroscience is how these dimensions are represented in the brain. Much neuroimaging work has examined valence and arousal separately and has identified partly dissociable neural correlates. Valence-related responses have been reported in the amygdala, insula, prefrontal cortex, and reward-related striatal regions, whereas arousal-related responses have been reported in the amygdala, thalamus, striatum, anterior cingulate cortex, and adjacent medial frontal cortex (Anders et al., 2004; Gerdes et al., 2010; Viinikainen et al., 2010). However, neither dimension alone fully characterizes the emotional content of a natural scene: images matched in valence can differ substantially in arousal, and images of opposite valence can evoke comparable levels of arousal. Determining how valence and arousal are jointly represented in the brain is therefore important for understanding the neural basis of visually evoked emotion.

Valence and arousal are not visual features in the same sense as luminance, contrast, color, or spatial layout. Instead, extracting the affective significance of a natural scene requires integrating visual information across multiple levels, including objects, actions, social interactions, and their relations within a scene (Bar, 2004; Bonner & Epstein, 2021; McMahon & Isik, 2023). We therefore predict that joint valence–arousal representations depend primarily on higher-level visual and semantic information and are distinct from representations of low-level visual features in both cortical location and temporal emergence. At this higher level, a scene is represented not only in terms of its elements, but also in terms of the narrative and semantics, how the elements relate to one another, and what the scene signifies for the observer. Its affective significance is one component of this broader scene description, with valence and arousal providing continuous dimensions of the scene’s pleasantness and intensity. Thus, neural representations of scene-level information should also contain information about valence and arousal (Abdel-Ghaffar et al., 2024). We further predict that regions representing joint valence and arousal will be contained within the more extensive network representing scene-level information and that scene-level representations will emerge before, or at least no later than, valence–arousal representations.

We tested these predictions by applying representational similarity analysis (RSA) to fMRI data and RSA-based EEG–fMRI fusion to simultaneously acquired EEG–fMRI data from 20 participants viewing 60 images from the International Affective Picture System (IAPS; Lang et al., 1997). We constructed three image-level representational dissimilarity matrices (RDMs): an Emotion RDM based on pairwise distances in the two-dimensional valence–arousal rating space, a GIST RDM based on pairwise distances in the space spanned by low-level visual features characterized by GIST descriptors, and a Scene RDM based on pairwise distances between CLIP-ViT image embeddings. Each image-based RDM was correlated with fMRI-based neural RDMs derived from pairwise distances between image-activated fMRI patterns in a whole-brain searchlight analysis and RSA-based EEG–fMRI fusion was then used to characterize the temporal dynamics of the corresponding neural representations. Finally, to assess whether the affective organization of scenes captured by human observers is also reflected in contemporary multimodal large language models (LLMs), we examined whether valence and arousal ratings generated by leading multimodal LLMs reproduced human ratings and predicted the joint neural representation of human-rated valence and arousal.

## Methods

### Participants

All procedures were approved by the Institutional Review Board of the University of Florida. Healthy volunteers (n = 26) with normal or corrected-to-normal vision participated in the study after providing written informed consent. They viewed images from the International Affective Picture System (IAPS) while simultaneous EEG–fMRI data were recorded. Two withdrew before completing data collection, and four were excluded for excessive head movement during scanning; data from these individuals were excluded. The analyzed sample comprised 20 participants (10 women; mean age ± SD: 20.4 ± 3.1 years).

### Experimental Procedure

#### The stimuli

The stimulus set comprised 60 grayscale IAPS pictures (Lang et al., 1997) with the following picture IDs: 4311, 4599, 4610, 4624, 4626, 4641, 4658, 4680, 4694, 4695, 2057, 2332, 2345, 8186, 8250, 2655, 4597, 4668, 4693, 8030, 2398, 2032, 2036, 2037, 2102, 2191, 2305, 2374, 2377, 2411, 2499, 2635, 2347, 5600, 5700, 5781, 5814, 5900, 8034, 2387, 1114, 1120, 1205, 1220, 1271, 1300, 1302, 1931, 3030, 3051, 3150, 6230, 6550, 9008, 9181, 9253, 9420, 9571, 3000, 3069. Each picture has associated normative valence and arousal ratings on 1–9 scales, where valence values of 1 and 9 indicate extremely unpleasant and extremely pleasant content, respectively, and arousal values of 1 and 9 indicate calm and highly arousing content, respectively.

#### Experimental paradigm

Participants completed five sessions of data collection (Figure 1B). In each session the 60 IAPS pictures were presented in random order. On each trial, a picture was displayed for 3 s on an MR-compatible monitor and viewed via a mirror mounted on the head coil, followed by a variable intertrial interval of 2,800 or 4,300 ms. Participants were asked to maintain fixation on a central fixation point throughout the recording session. Across the whole experiment each picture was repeated five times for a total of 5 × 60 = 300 trials.

**Figure 1.**
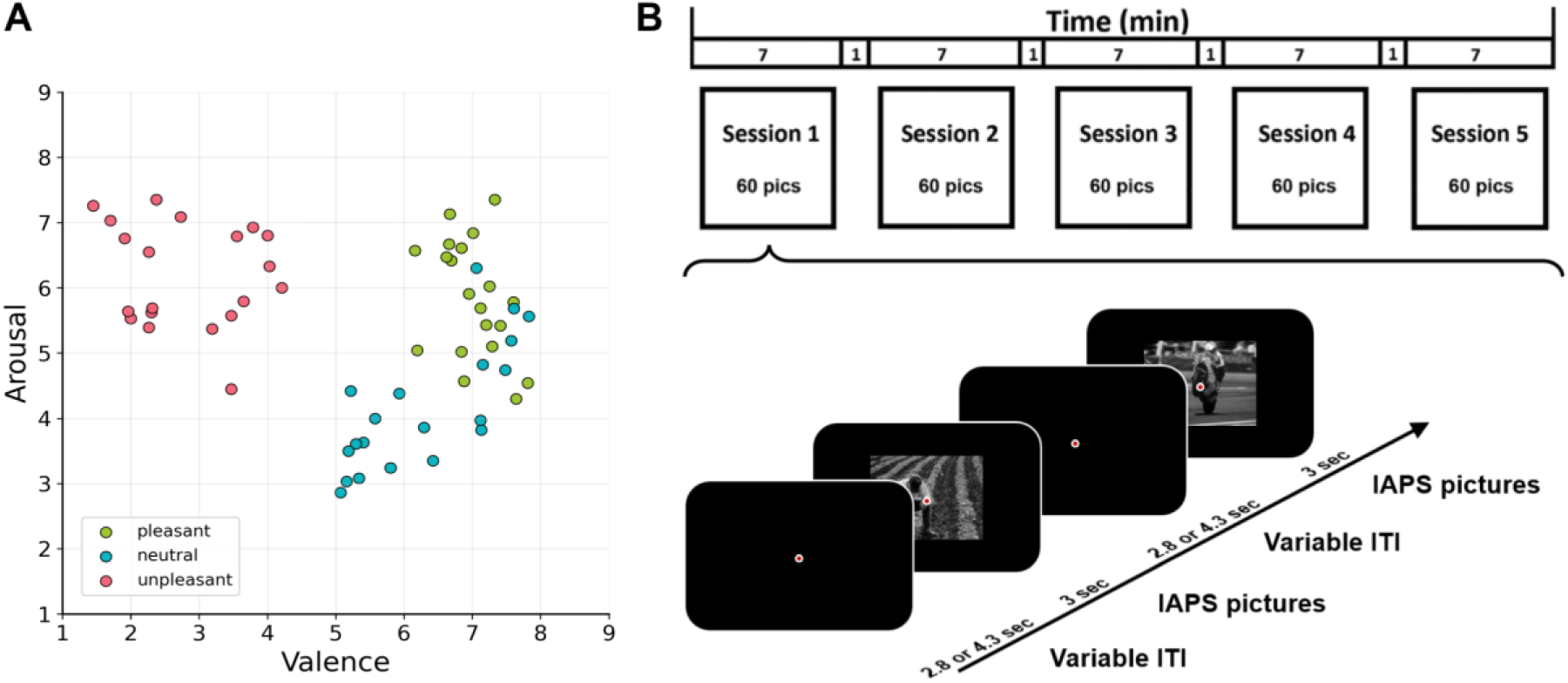
Stimuli and experiment design. **(A)** 60 IAPS images represented on the valence and arousal plane. These images can be broadly labeled as pleasant, unpleasant or neutral. **(B)** The experiment consisted of five passive-viewing sessions. The same 60 IAPS images were presented in each session with the presentation order randomized from session to session. The timeline of a representative trial is illustrated. Participants were asked to maintain fixation during picture presentations. ITI: Intertrial interval.

### Data Acquisition and Preprocessing

#### fMRI data

Functional MRI (fMRI) data were collected on a 3T Philips Achieva scanner (Philips Medical Systems) with the following parameters: echo time (TE) = 30 ms; repetition time (TR) = 1.98 s; flip angle = 80°; 36 slices; field of view = 224 mm; voxel size = 3.5 × 3.5 × 3.5 mm; matrix size = 64 × 64. Slices were acquired in ascending order and oriented parallel to the plane connecting the anterior and posterior commissures. High-resolution T1-weighted structural images were also obtained for anatomical reference and normalization.

The fMRI data were preprocessed using SPM (http://www.fil.ion.ucl.ac.uk/spm/). The first five volumes from each session were discarded to eliminate artifacts associated with transient scanner instability. Slice timing correction was applied using interpolation to account for differences in slice acquisition time. Images were then corrected for head movement by spatial realignment to the sixth image of each session, normalized to the Montreal Neurological Institute (MNI) template, and resampled to 3 × 3 × 3 mm spatial resolution. The transformed images were smoothed using a Gaussian filter with 8 mm full width at half maximum (FWHM). Low-frequency temporal drift was removed using a high-pass filter with a cutoff period of 128 s.

#### EEG data

EEG data were recorded simultaneously with fMRI using a 32-channel MR-compatible EEG system (Brain Products GmbH). Thirty-one sintered Ag/AgCl electrodes were placed on the scalp according to the 10-20 system, with FCz serving as the reference electrode. An additional electrode was placed on the subject’s upper back to monitor electrocardiogram (ECG) for detecting heartbeat events and assisting in cardioballistic artifact removal during preprocessing. EEG signals were recorded with an online 0.1-250 Hz band-pass filter and digitized to 16-bit at a sampling rate of 5 kHz. The EEG recording system was synchronized with the scanner’s internal clock throughout recording to ensure successful gradient artifact removal in subsequent analyses.

The EEG data were first preprocessed using Brain Vision Analyzer 2.0 (Brain Products GmbH, Germany) to remove scanner-induced artifacts. Gradient artifacts were removed using a modified average artifact subtraction method (Allen et al., 2000): an artifact template was created by segmenting and averaging the data according to the onset of each MR volume acquisition and subtracted from the raw EEG data. Cardioballistic artifacts were removed using an average artifact subtraction method (Allen et al., 1998): R peaks were detected in the low-pass-filtered ECG signal and used to construct a delayed average artifact template over 21 consecutive heartbeat events using a sliding-window approach, which was then subtracted from the original signal. After gradient and cardioballistic artifact removal, the EEG data were low-pass filtered at 50 Hz, downsampled to 250 Hz, re-referenced to the average reference, and exported to EEGLAB (Delorme & Makeig, 2004) for further processing. Second-order blind identification (SOBI; Belouchrani et al., 1997) was applied to correct eye blinking, residual cardioballistic artifacts, and movement-related artifacts. The artifact-corrected data were then low-pass filtered at 30 Hz and epoched from −300 to 2000 ms relative to picture onset (0 ms).

### Single-trial Estimation of fMRI Response

All analyses were done at the single trial level. Trial-by-trial picture-evoked BOLD signals were estimated using the beta series method (Mumford et al., 2012). In this approach, a trial of interest was modeled by one regressor while all remaining trials were modeled by another regressor. Six motion parameters were included as additional regressors to account for movement-related artifacts. This procedure was repeated for all trials, yielding single-trial BOLD response estimates (beta values) in every voxel for each picture presentation. These single-trial BOLD responses served as input for the RSA analysis.

### Representational Similarity Analysis (RSA)

#### Neural representation of valence–arousal

RSA analysis was applied to examine the joint neural representation of valence and arousal (Kriegeskorte et al., 2008). It consists of constructing a representational dissimilarity matrix (RDM) for 60 IAPS images and a RDM for the neural activity in a region of interest (ROI) evoked by the 60 images and correlating the two RDMs. Specifically, for any pair of IAPS images, their valence–arousal dissimilarity was defined as the Euclidean distance between the two images on the valence–arousal plane (Figure 2A), whereas their neural dissimilarity was measured by the Euclidean distance between the voxel-level fMRI responses evoked by the two images; here, the larger the distance, the more dissimilar the two images. Euclidean distance measures, derived from valence/arousal space, have been used previously to jointly describe a picture’s motivational direction and intensity, sometimes interpreted as the strength of appetitive and aversive motivation (Keil and Freund, 2009). A brain region with higher RSA correlation between the two RDMs is considered more strongly involved in the joint representation of valence and arousal. Hereafter, we refer to the valence–arousal-based RDM as the Emotion RDM.

**Figure 2.**
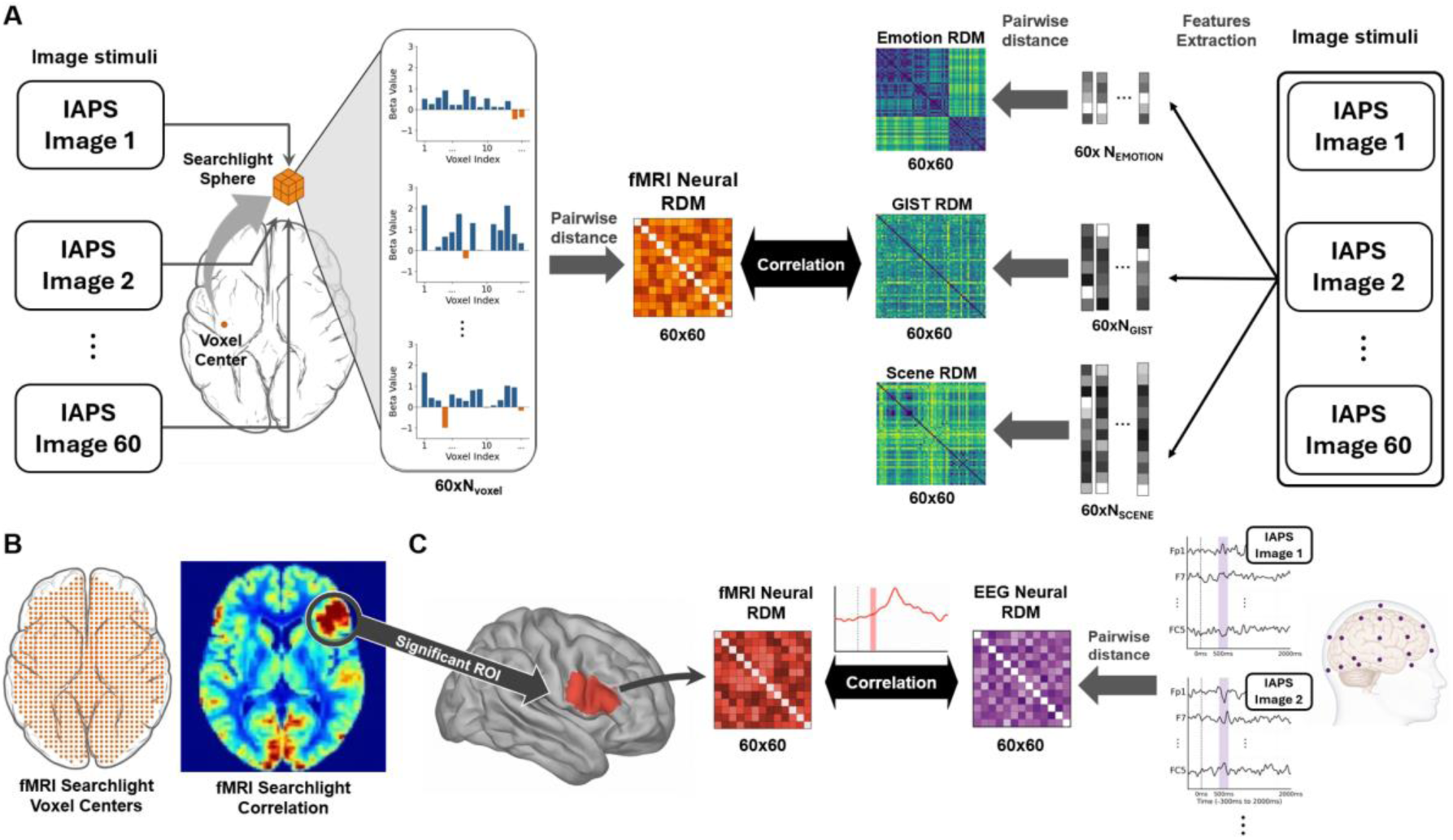
Overview of the searchlight fMRI RSA and EEG–fMRI fusion analysis framework. **(A)** Neural RDMs were derived from local fMRI response patterns and correlated with Emotion RDM, GIST RDM and Scene RDM. **(B)** Whole-brain searchlight analysis yielded brain maps revealing the neural representations of valence–arousal, low-level visual properties, and scene-level properties. **(C)** EEG–fMRI fusion uncovered the temporal dynamics of valence-arousal, low-level visual and scene-level visual representations.

#### Neural representation of low-level visual features

For comparison, we also assessed the neural representation of low-level visual features. Each grayscale image was passed through a set of Gabor filters at multiple orientations and spatial frequencies in a 4 × 4 grid, yielding a 512-dimensional feature vector, called the GIST vector, which summarizes the image’s “spatial envelope” (Oliva & Torralba, 2001). Pairwise Euclidean distances between all pairs of the 60 GIST vectors formed a 60 × 60 RDM reflecting the dissimilarity structure of the low-level visual features of the IAPS pictures (Figure 2A). The GIST RDM was correlated with the neural RDM described above. A brain region with higher RSA correlation is considered more strongly involved in the representation of low-level visual features. Hereafter, we refer to the low-level visual feature RDM as the GIST RDM.

#### Neural representation of scene-level features

To model scene-level representation, we extracted high-level image embeddings from CLIP-ViT-B/32, a multimodal vision-language model trained on hundreds of millions of image-text pairs (Radford et al., 2021). For each of the 60 images, we extracted a 512-dimensional embedding vector from the final projection layer of the model’s image encoder. These embeddings capture scene-level properties, including object identity, spatial layout, contextual structure, and semantic content. Pairwise Euclidean distances between the 512-dimensional CLIP-ViT image embeddings yielded a 60 × 60 scene-level representational dissimilarity matrix (Figure 2A). This scene-level RDM was then correlated with the neural RDM described above. A brain region with higher RSA correlation is considered more strongly involved in scene representation. Hereafter, we refer to this scene-level RDM as the Scene RDM.

#### Searchlight RSA

To map the representation of stimulus properties over the whole brain, we conducted a searchlight RSA analysis using the fMRI data (Kriegeskorte et al., 2006). For each gray-matter voxel, we defined a 15-mm-radius sphere centered on the voxel, containing approximately 484 in-mask voxels; see Supplementary Figure S1 for analysis underlying the choice of the 15 mm radius for the searchlight sphere. A 60 × 60 local neural RDM for each sphere was constructed using correlation distance (1 − Pearson’s *r*) between the response patterns evoked by every pair of images. All neural and model RDMs were min-max normalized to the range [0, 1]. The upper-triangular elements of the neural RDM were then correlated with the Emotion RDM, the GIST RDM, and the Scene RDM, yielding one correlation value per model at each voxel center (Figure 2A). This procedure was repeated across all gray-matter voxels to generate participant-level whole-brain correlation maps. The correlation values were Fisher-transformed and entered into group-level one-sample *t*-tests (two-sided, retaining positive t values). Voxelwise *p* values were adjusted using the Benjamini-Hochberg FDR procedure and thresholded at FDR-corrected p < .05. Clusters containing at least 50 such voxels were retained (Figure 2B). ROIs so identified were used for EEG–fMRI fusion analysis to examine the temporal emergence of the neural representation (see below).

#### Cortical atlas annotation

Functional maps were displayed on fsaverage cortical flatmaps using pycortex (Gao et al., 2015) and annotated with 16 anatomical regions derived from HCP-MMP1.0 parcels (Glasser et al., 2016). To describe the lateral visual and temporo-parietal anatomy consistently across maps, we defined the middle temporal complex (MT+) as the union of MT, MST, FST, and V4t; lateral occipital cortex (LOC) as LO1, LO2, and LO3; and the temporo-parietal junction (TPJ) as TPOJ1, TPOJ2, and TPOJ3. The remaining group definitions and voxel-assignment rule are provided in Supplementary Table S1. Atlas outlines indicate anatomical extents, not the extent or significance of functional effects.

#### EEG–fMRI fusion

EEG and fMRI were fused to examine the temporal dynamics of the neural representation of given stimulus properties. For each representation, an ROI was defined as the set of voxels surviving thresholding in its group-level fMRI searchlight map; see above. Within each ROI, a 60 × 60 fMRI RDM was calculated using pairwise Euclidean distances between fMRI patterns evoked by the 60 images. For EEG, the five repetitions of each image were averaged, and a 60 × 60 RDM was calculated from the 31-channel activity pattern at every 4-ms sample from −300 to 2000 ms using the Euclidean distance measure. At each time point, the upper-triangular elements of the EEG RDM were correlated with those of the corresponding ROI-level fMRI RDM (Cichy et al., 2014), producing a participant-level fusion time course for each representation map (Figure 2C). Each participant’s fusion time course was baseline-corrected, averaged across participants, and smoothed with a Gaussian kernel with a standard deviation of 10 samples (40 ms). The emergence time of each representation was defined as the first time point at which the smoothed fusion time course was positive and exceeded five standard deviations of baseline fluctuations for at least 50 consecutive 4-ms samples (200 ms).

For the representations of the three stimulus properties, the differences in their emergence onset times were assessed using a bootstrap resampling procedure (Bo et al., 2022). Participant-level EEG–fMRI RSA time courses were resampled with replacement 10,000 times. For each resample, the group-average EEG–fMRI fusion time course was recomputed and the emergence onset was defined using the same method described above. The three bootstrap distributions of emergence onset times were compared pairwise using two-sided two-sample Kolmogorov-Smirnov tests.

### Univariate Analysis of Joint Valence–Arousal Representation

While the main focus here is on the joint multivariate representation of valence and arousal, we also performed a complementary univariate analysis, examining whether at the single voxel level, picture-evoked BOLD responses were jointly predicted by valence and arousal ratings. For each participant, a multiple regression was carried out independently at every voxel, with valence and arousal ratings entered simultaneously as predictor variables and the picture-evoked BOLD response as the dependent variable. The valence coefficient therefore reflected the association between valence and the BOLD response after accounting for arousal, and the arousal coefficient reflected the association between arousal and the BOLD response after accounting for valence. At the group level, participant-specific valence and arousal coefficients were tested against zero using separate two-sided one-sample t-tests. The resulting voxelwise *p* values were adjusted separately for the valence and arousal maps using the Benjamini-Hochberg FDR procedure (FDR-corrected p < .05). Positive and negative effects were cluster-thresholded separately, and clusters containing at least 50 voxels were retained.

To identify regions showing positive effects of both affective dimensions, we intersected the positive valence and arousal maps after FDR and cluster-extent thresholding. For visualization, each voxel in this conjunction map was assigned the smaller of its valence and arousal *t* values, such that the displayed value reflected the weaker of the two effects. The minimum t value was used only for visualization; inclusion in the conjunction required significant positive coefficients for both predictors. No additional cluster-extent threshold was applied after the conjunction was formed.

### Neural Representations of LLM-Based Valence and Arousal Ratings

Valence and arousal ratings for affective visual materials, including pictures and videos, have traditionally been obtained from human raters. This process can be time-consuming and costly, and rating quality can vary greatly across raters. We tested the extent to which multimodal large language models (LLMs) can be used to rate valence and arousal. Three LLMs were used for this purpose: ChatGPT o3, Claude 4 Sonnet, and Gemini 2.5 Pro. Each model was given the 60 IAPS images and asked to provide valence and arousal ratings on a 1–9 scale. Each image was rated using three prompts, and the resulting ratings were averaged. See the Supplementary Materials for examples of the prompts used in this study.

Similar to human ratings, AI-generated valence and arousal ratings were treated as coordinates on a two-dimensional valence–arousal plane, and an AI Emotion RDM was derived for each model by computing pairwise Euclidean distances between images. We first quantified representational correspondence between AI and human ratings by correlating each AI Emotion RDM with the human Emotion RDM using Spearman correlation. We then assessed neural representations corresponding to the AI-derived Emotion RDMs using the fMRI searchlight RSA analysis described above. The spatial overlap between the human and AI valence–arousal representations was quantified using the Jaccard index, defined as the number of overlapping voxels divided by the total number of voxels in the union of the two maps. For Figure 6C–E, we retained the 1,038 voxels with the largest positive t values from each FDR p < .05, cluster-filtered map to match the human map size. At each retention level from 100% to 10% in 10% steps, we selected the highest-ranked voxels within each matched map. To construct the null distribution, we independently drew the same number of voxels for each map without replacement from the 59,159 eligible voxel centers in the searchlight analysis mask, repeating this procedure 1,000 times at each retention level. One-sided empirical p values were calculated as (1 + the number of null Jaccard values at least as large as observed) / 1,001. This random-set null tests spatial coincidence under independent voxel placement within the analysis mask and does not preserve the clustering of the observed maps.

## Results

### Characterization of Image Properties

The stimulus set consisted of 60 IAPS images. Each image was characterized using three sets of properties: affective properties, indexed by valence and arousal ratings; low-level visual properties, indexed by GIST descriptors; and scene-level visual properties, indexed by CLIP-ViT image embeddings. For each set of properties, we quantified the similarity structure among images using a representational dissimilarity matrix (RDM). As shown in Figure 3A, the Emotion RDM was significantly correlated with the Scene RDM (*r* ≈ 0.29, *p* = 3.15 × 10^−39^), but not with the GIST RDM (*r* ≈ −0.04, *p* > .05). The Scene RDM was likewise not significantly correlated with the GIST RDM (*r* ≈ −0.02, *p* > .05). These findings indicate that the affective similarity structure of the images was related to their scene-level semantic similarity structure, whereas neither was related to the similarity structure derived from low-level visual properties.

**Figure 3.**
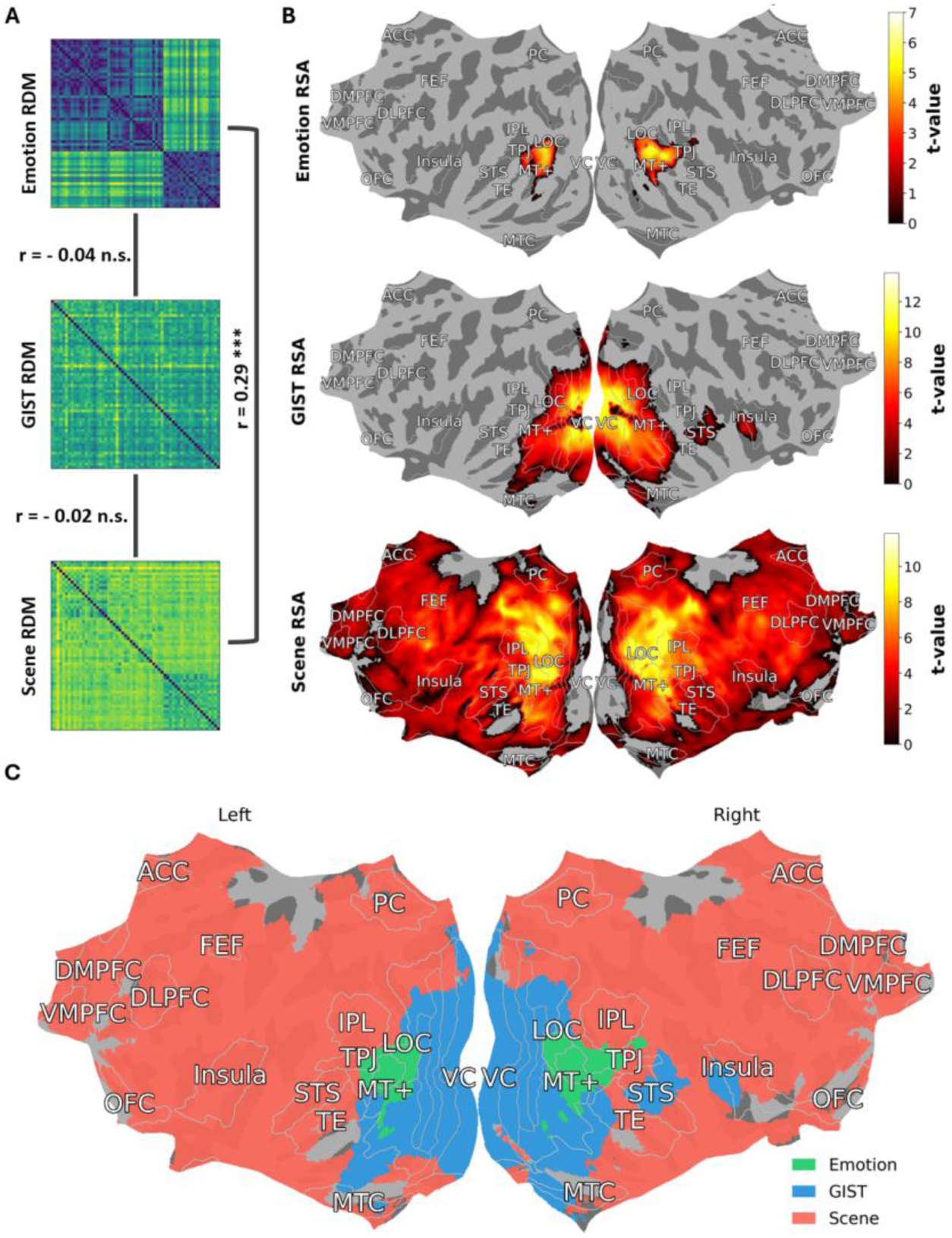
Image properties and their multivariate neural representations. **(A)** Pairwise correlation of the three RDMs. The GIST RDM was not significantly correlated with either the Emotion RDM or the Scene RDM, whereas the Emotion and Scene RDMs showed a significant positive correlation. **(B)** fMRI searchlight analysis. The Emotion RDM is significantly correlated with neural RDM mainly in MT+, LOC, TPJ, and higher-order VC (top); the GIST RDM was significantly correlated with neural RDMs primarily in visual cortex, especially early visual areas such as V1, with additional involvement of the right STS and right insula (middle); and the Scene RDM is significantly correlated with neural RDM in broadly distributed areas in occipital, temporal, parietal, and frontal cortices (bottom). All maps were thresholded at FDR-corrected p < .05 with the additional requirement that the cluster size is at least 50 voxels. **(C)** Overlap of the three maps in (B): the Emotion map (green), GIST map (blue), and Scene map (red).

### Neural Representations of Image Properties: Multivariate Analysis

Applying searchlight RSA to the fMRI data, we identified brain regions in which local neural RDMs were significantly correlated with each of the three model RDMs (Figure 3B). Representation maps were thresholded at FDR-corrected p < .05, with an additional cluster-extent threshold of at least 50 voxels.

The Emotion RDM showed significant positive correlations primarily in bilateral MT+, LOC, and TPJ, with additional areas in VC (Figure 3B, top). The GIST RDM showed significant positive correlations in retinotopic visual areas V1, V2, V3, and V4, as well as higher-order visual areas including fusiform face complex (FFC), ventral visual complex (VVC), LOC, TE, MT+, and MTC, with additional involvement of right STS and right insula (Figure 3B, middle). The Scene RDM showed significant positive correlations across broadly distributed regions in occipital, temporal, parietal, and frontal cortex, including VC, LOC, MT+, TPJ, STS, FEF, insula, ACC, mPFC, and PC (Figure 3B, bottom).

The valence-arousal map had 1,038 voxels and was fully contained within the scene representation map (41,812 voxels). The GIST representation map had 7,921 voxels, 538 of which also appeared in the Emotion map, where the overlap, corresponding to 51.8% of the Emotion map and 6.8% of the GIST map, was located primarily in bilateral inferior lateral occipital cortex, with smaller extensions into temporo-occipital fusiform and inferior temporal cortex. The Scene and GIST maps showed a larger overlap of 5,379 voxels, corresponding to 67.9% of the GIST map and 12.9% of the much broader Scene map. The overlap was concentrated bilaterally in the inferior lateral occipital cortex and occipital pole and extended into occipital and temporo-occipital fusiform cortex and adjacent temporo-occipital inferior temporal cortex. This partial convergence is consistent with the features represented by the two models: GIST characterizes orientation, spatial frequency, texture, and coarse spatial layout, whereas CLIP-ViT captures higher-level scene properties including semantic content while retaining information about lower-level features such as visual form and spatial layout.

Because the Emotion map was fully contained within the Scene map and the two model RDMs were significantly correlated, spatial overlap alone could not determine whether neural representational structure within the Emotion-defined region reflected valence-arousal relationships, broader scene relationships, or both. We therefore regressed the neural RDMs against the Emotion and Scene RDMs simultaneously and partitioned the explained variance into contributions unique to each model and a component shared by both models. Here, unique variance refers to the variance explained by one RDM after accounting for the other RDM. The combined model explained R² = 0.0105 amount of variance. The Emotion RDM explained variance was R² = 0.0029 (FDR-corrected *p* = 1.12 × 10⁻¹¹), whereas the Scene RDM explained variance was R² = 0.0086 (FDR-corrected *p* = 1.25 × 10⁻⁹). The component attributable to variance shared by the two predictors was not significant (R² = −0.0010; FDR-corrected *p* > 0.05). Thus, neural similarity within the Emotion-defined region contained separable valence-arousal and scene-related components rather than being explained solely by information common to the two model RDMs.

### Temporal Dynamics of Neural Representations

EEG and fMRI data were fused using RSA to examine the temporal dynamics of neural representations of the three sets of image properties. EEG RDMs were constructed at each time point in the EEG time series and correlated with fMRI RDMs derived from the three representation maps identified in Figure 3B. Specifically, for each set of image properties, the entire representation map was treated as a region of interest (ROI), from which the fMRI RDM was derived. As shown in Figure 4A, the GIST-defined time course increased rapidly after stimulus onset, becoming significant at approximately 124 ms post-stimulus onset and remaining above threshold throughout the analyzed time window. This pattern suggests early and sustained representation of low-level visual features. In contrast, the Scene-defined and Emotion-defined time courses remained near baseline during the early post-stimulus period and reached significance later, at approximately 428 ms and 532 ms, respectively. To compare statistically the emergence onset times across the three EEG–fMRI fusion time courses, we conducted a bootstrap procedure. The resulting bootstrap distributions of emergence onset times are shown in Figure 4B. Pairwise two-sided Kolmogorov–Smirnov tests indicated that all three onset-time distributions differed significantly from one another (*p* < .001). Together, these findings suggest the following temporal sequence of representational emergence: low-level visual properties at 124 ms, followed by scene-level properties at 428 ms, and then affective properties at 532 ms.

**Figure 4.**
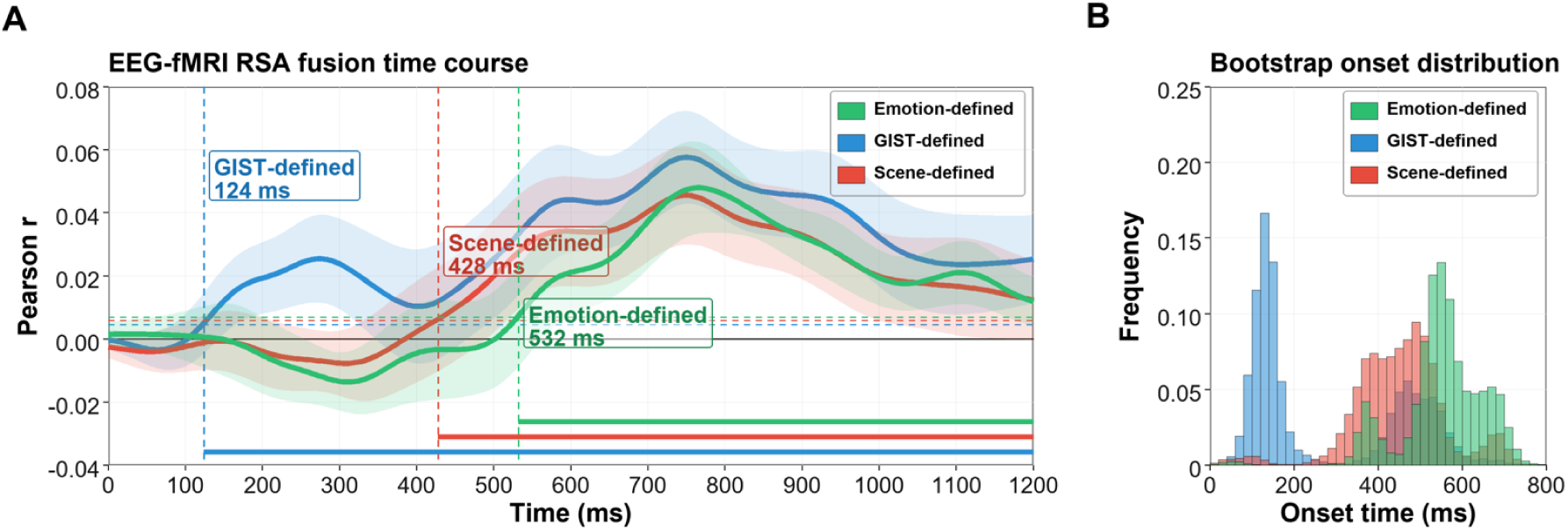
Temporal dynamics of neural representations. **(A)** EEG–fMRI fusion time courses. EEG RDMs at every time point were correlated with neural RDMs from each of the three maps: Emotion, GIST, and Scene maps to reveal the temporal emergence of the neural representations of the three sets of image properties. Here shading reflects SEM, dotted horizontal lines show representation-specific five-SD thresholds, dashed vertical lines mark the onset times, and solid horizontal lines show threshold-exceeding intervals. **(B)** Bootstrap distributions of emergence onset times for the three time courses in (A).

### Neural Representation of Valence and Arousal: Univariate fMRI Analysis

To complement the multivariate analysis, we examined univariate representations of valence and arousal using multiple regression and compared the resulting map with the multivariate representation. The univariate map comprised 419 voxels, of which 234 overlapped with the multivariate map (Figure 5A). Thus, the overlapping constituted 55.8% of the univariate map and 22.5% of the multivariate map. Spatial convergence was concentrated primarily in bilateral occipitotemporal cortex. Both maps contained voxels in bilateral LOC, MT+, and TPJ, as well as right VC. The multivariate map additionally included the left VC and TE, whereas the univariate map additionally included the left DMPFC and VMPFC, and bilateral ACC. Figure 5B lists regions common to both representations, as well as regions specific to the multivariate or univariate representation.

**Figure 5.**
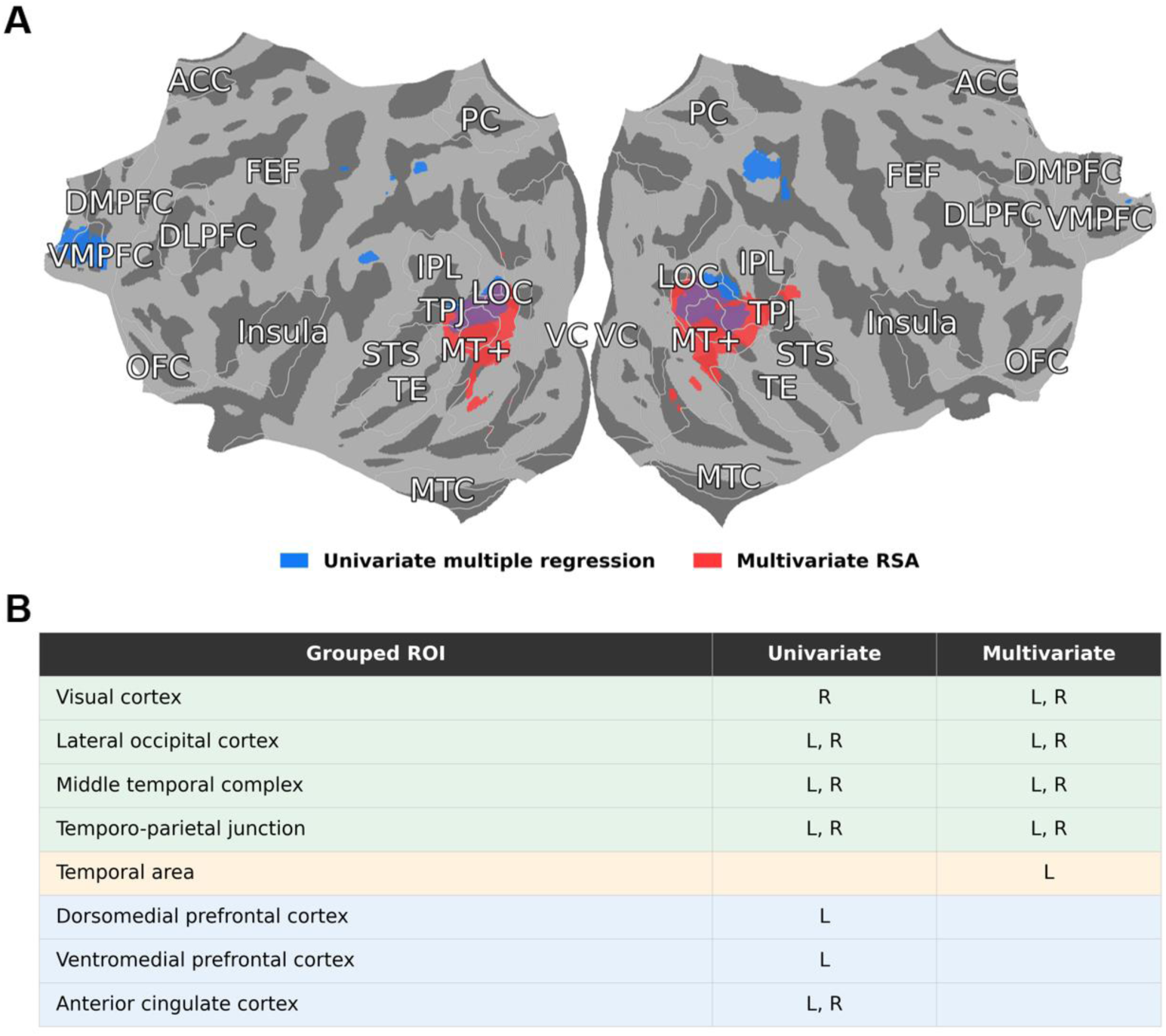
Univariate analysis of valence–arousal representation. **(A)** Univariate (blue) representation map superimposed on the multivariate representation map (red) from Figure 3B. **(B)** Anatomical locations of the two maps. The univariate conjunction uses FDR-corrected p < .05 for each predictor before intersection; the multivariate map uses FDR-corrected p < .05.

### Neural Representations of LLM-based Valence and Arousal

Valence and arousal ratings were obtained from three AI models (ChatGPT o3, Claude 4 Sonnet, and Gemini 2.5 Pro) for the 60 IAPS images. Figure 6A displays the AI-generated ratings together with human ratings in the valence–arousal space, and Figure 6B shows pairwise comparisons among the four Emotion RDMs. Spearman correlations between each AI Emotion RDM and the human Emotion RDM were strong: ChatGPT o3, *r* = 0.79; Gemini 2.5 Pro, *r* = 0.72; and Claude 4 Sonnet, *r* = 0.70; all *p* < .001. The three AI Emotion RDMs were also strongly correlated with one another (mean AI–AI correlation, *r* = 0.88), where the mean correlation exceeded their mean correlation with the human Emotion RDM (mean human–AI correlation, *r* = 0.73). In the fMRI searchlight RSA analysis, neural representation maps derived from each AI Emotion RDM largely overlapped with the human valence–arousal representation map (Figure 6C). For all three AI models, peak correlations were located in lateral occipitotemporal cortex, in agreement with the human map. The Gemini 2.5 Pro map additionally included a cluster in calcarine cortex that was not observed for ChatGPT o3 or Claude 4 Sonnet and was also not in the human map. All maps were thresholded at FDR-corrected p < .05, with a cluster-extent threshold of at least 50 voxels.

**Figure 6.**
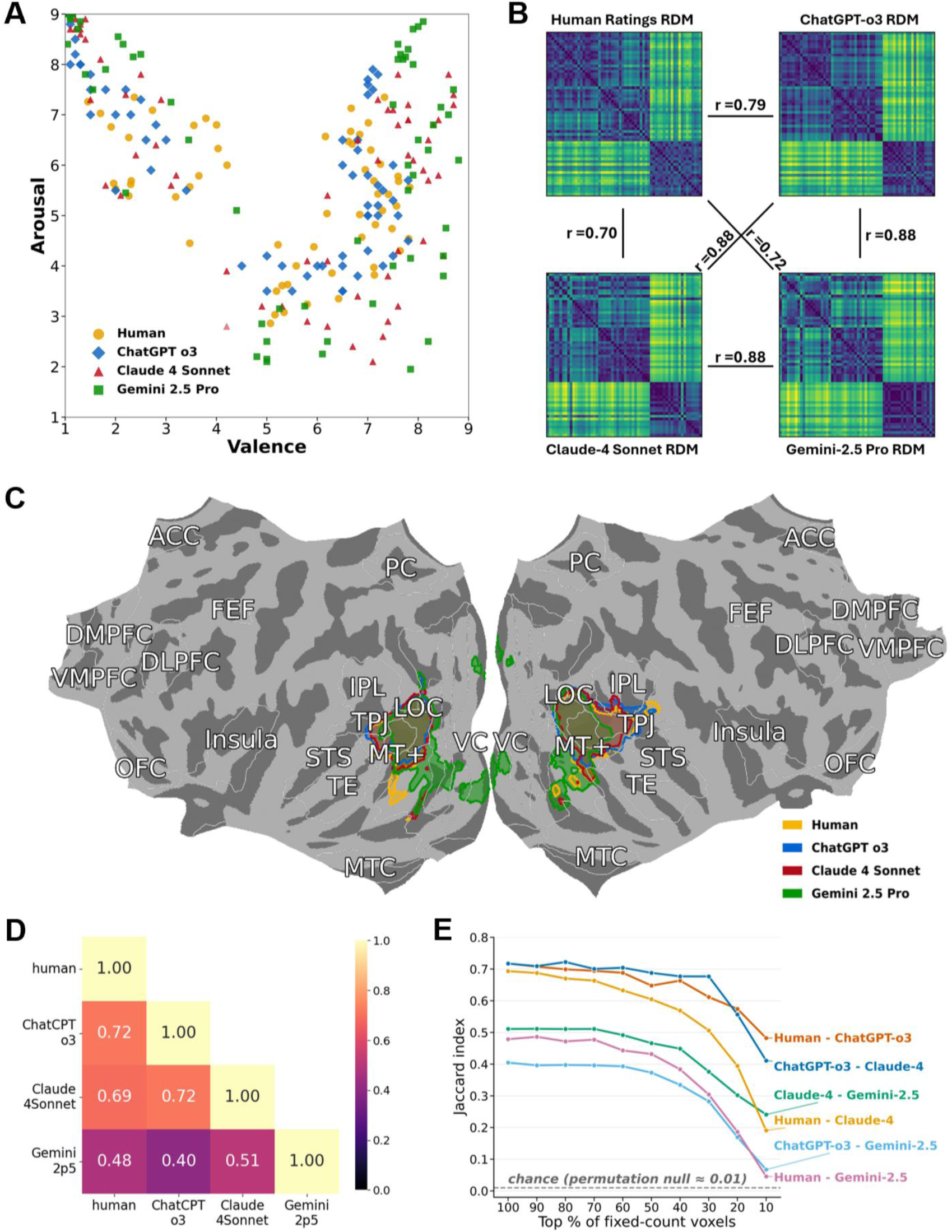
AI-generated valence and arousal and their joint neural representations. **(A)** AI-generated valence and arousal ratings and human ratings show similar patterns in the valence-arousal space. **(B)** The four Emotional RDMs (human RDM and three AI RDMs) exhibited strong pairwise correlations. **(C)** RSA searchlight analysis shows highly overlapped joint valence–arousal representation maps. **(D)** Jaccard indices for pairwise spatial overlap of representation maps in (C). **(E)** Pairwise Jaccard indices as a function of voxel-retention threshold (colored lines). The gray dashed curve shows the mean Jaccard index from 1,000 random-set permutations at each threshold.

We used the Jaccard index to quantify spatial overlap. Voxel counts were matched across the four maps by retaining the top 1,038 voxels ranked according to the t values in each map. The human map was the smallest. Before matching, the full thresholded maps contained 1,038 human, 1,105 ChatGPT o3, 1,156 Claude 4 Sonnet, and 2,658 Gemini 2.5 Pro voxels. As shown in Figure 6D, human–AI overlap was highest for ChatGPT o3 (*J* = 0.72), followed by Claude 4 Sonnet (*J* = 0.69) and Gemini 2.5 Pro (*J* = 0.48). Overlap among the AI maps was of a similar magnitude: ChatGPT o3 versus Claude 4 Sonnet, *J* = 0.72; Claude 4 Sonnet versus Gemini 2.5 Pro, *J* = 0.51; and ChatGPT o3 versus Gemini 2.5 Pro, *J* = 0.40. Figure 6E shows how the observed pairwise overlaps varied as progressively fewer voxels were retained. Overlap declined with more selective voxel sets, particularly for Gemini 2.5 Pro. For the permutation benchmark, we independently drew equally sized voxel sets from the 59,159-voxel searchlight analysis mask in 1,000 iterations at each retention level (Figure 6E). At the full 1,038-voxel level, the mean null Jaccard index was approximately 0.009, and none of the null draws reached an observed human–AI overlap (all one-sided empirical p < .001): human–ChatGPT o3, z = 343.2; human–Claude 4 Sonnet, z = 334.6; and human–Gemini 2.5 Pro, z = 229.7. All six pairwise overlaps exceeded every null draw at each tested retention level from 100% through 10% (all empirical p < .001). Together, these results indicate robust spatial agreement between representations of AI-derived and human-derived valence and arousal across relatively broad spatial scales.

## Discussion

In this study, we obtained the joint neural representation of valence and arousal and compared it with representations of low-level and scene-level visual properties. Searchlight RSA of the fMRI data showed that the joint valence–arousal representation was localized primarily in higher-order visual areas including MT, MST, LOC, and TPJ. This representation showed limited spatial overlap with the representation of low-level visual properties, which was centered primarily in early visual cortex, but was entirely contained within the more broadly distributed representation of scene-level visual properties. EEG–fMRI fusion further indicated that the representation of low-level visual properties emerged at approximately 124 ms after stimulus onset, followed by the scene representation at approximately 428 ms, and the valence–arousal representation at approximately 532 ms. Finally, the valence and arousal ratings generated by three multimodal LLMs showed close correspondence with human valence and arousal ratings, and the neural representation maps derived from these LLM-based ratings had substantial spatial overlap with that derived from human valence and arousal ratings.

### Joint Representation of Valence and Arousal

Valence and arousal are widely regarded as fundamental dimensions of affect (Russell and Barrett, 1999; Lang and Bradley, 2010). Valence describes the extent to which a stimulus is experienced as pleasant or unpleasant, whereas arousal reflects the degree of activation or intensity it elicits. Together, these dimensions provide a compact framework for characterizing self-reported affect across a broad range of stimuli. For natural scenes, however, neither dimension alone adequately captures their affective organization. Scenes with similar arousal levels can differ markedly in valence: an exciting sporting event and a threatening attack may both be highly arousing but evoke opposite affective evaluations. Likewise, scenes with similar valence can differ in arousal: a serene landscape and an erotic image may both be positive but differ substantially in intensity and motivational relevance. Considering valence and arousal jointly therefore preserves distinctions among affective states that would be obscured by either dimension alone (Bradley & Lang, 1994; Lang & Bradley, 2010).

Our results indicate that the joint valence–arousal structure was represented in bilateral lateral occipitotemporal and temporoparietal cortex, including MT, MST, LOC, and TPJ. Prior work links these regions to visual, semantic, and social interpretation of scenes. For example, multivoxel patterns across visual cortex, including LOC, distinguish affective from neutral scenes (Bo et al., 2021). In a functionally localized LOC, neural patterns also predict human emotion ratings for everyday scenes (Gao et al., 2025), consistent with the possibility that object representations contribute information relevant to affective appraisal. In functionally localized TPJ, neural patterns distinguish emotions inferred from described situations and reflect abstract event features (Skerry & Saxe, 2015). Importantly, those representations extended beyond valence and arousal, linking emotion attribution to conceptual knowledge about events rather than establishing a valence–arousal code per se.

The involvement of MT and MST is also notable. These regions are strongly associated with visual-motion processing, and affective content in static scenes can depend on implied motion or action, such as a person fleeing danger, an approaching threat, or an opportunity for approach. MT is a central component of the proposed third visual pathway for social perception, which links early visual cortex through MT/V5 to posterior STS and has been proposed to support processing of dynamic social cues (Pitcher et al., 2014; Pitcher & Ungerleider, 2021). Although STS did not meet the threshold for inclusion in the valence–arousal map, the functional-connectivity analysis shown in Figure S2 indicates that it was functionally connected to regions within the map.

### Comparison with Representations of Low-level and Scene-level Visual Properties

We compared the joint valence–arousal representation with representations of low-level and scene-level visual properties. Low-level visual properties were modeled using the GIST descriptor, which summarizes an image’s orientations, spatial frequencies, and spatial layout (Oliva & Torralba, 2001). Scene-level visual properties were modeled using CLIP-ViT-B/32 embeddings. Unlike a hand-engineered low-level descriptor or a model trained solely for object classification, CLIP-ViT-B/32 is trained on large-scale image–text pairs. Its image embeddings are therefore shaped by distinctions relevant to how people describe scenes, including object identity, spatial context, actions, social content, and broader scene meaning (Radford et al., 2021). These embeddings thus provided a semantically rich model for assessing whether joint valence–arousal structure reflects low-level image statistics, scene-level semantic structure, or a related but distinct form of affective organization.

The similarity structures of the three RDMs provided initial evidence for this distinction. The Emotion RDM was not significantly associated with the GIST RDM, indicating that affective similarity among the IAPS images was largely independent of similarity in their low-level visual properties captured by GIST. In contrast, the Emotion RDM showed a moderate but significant association with the Scene RDM. Thus, the valence–arousal structure shared some organization with scene-level semantic content, while remaining distinct from both scene-level semantics and low-level image properties. This pattern is consistent with the view that affective meaning is partly grounded in scene content while also organizing images according to their evaluative significance.

The fMRI RSA maps revealed a corresponding spatial dissociation in neural representation. The representation of low-level visual properties was concentrated in early visual cortex, including the occipital pole, calcarine sulcus, and retinotopic visual areas, with smaller extensions into right STS and right insula. This map showed limited overlap with the valence–arousal representation, which was localized primarily in bilateral lateral occipitotemporal and temporoparietal cortex, including MT, MST, LOC, and TPJ. In contrast, the scene-level representation produced the most spatially extensive pattern, spanning broad temporal, parietal and frontal cortical regions, with additional occipital involvement. The emotion-defined map was entirely contained within this broader scene-defined map. This nesting suggests that the joint valence–arousal information, which may be interpreted as appetitive and aversive motivational intensity evoked by a given picture, involves specific activation patterns within a subset or regions that make up the network involved in encoding semantic scene content.

The temporal dynamics of representational emergence were consistent with this spatial organization. EEG–fMRI fusion showed that the representation of low-level visual properties emerged first, at approximately 124 ms after picture onset, followed by scene and valence–arousal representations at approximately 428 ms and 532 ms, respectively. The emergence of the valence–arousal representation during this later interval overlaps temporally with the late positive potential, a sustained ERP component associated with attention to and evaluative processing of motivationally significant stimuli (Cuthbert et al., 2000; Schupp et al., 2006; Hajcak et al., 2010). The lateral occipitotemporal locus of the valence–arousal representation is also consistent with prior evidence that inferotemporal and lateral occipitotemporal cortex encode affective dimensions across visual stimuli (Sabatinelli et al., 2005; Sabatinelli et al., 2009; Kragel et al., 2019), as well as more recent evidence for joint semantic and affective tuning in occipitotemporal cortex (Abdel-Ghaffar et al., 2024).

### Structure and Representation of AI-generated Valence and Arousal

The capabilities of large language models (LLMs) have advanced rapidly, making them a useful tool for testing whether they can be used to estimate the affective meaning of natural scenes. In the present study, LLMs were not treated as models of subjective emotional experience. Rather, they generated valence and arousal ratings for natural scenes, which were used as semantically informed estimates of how those scenes are likely to be evaluated. This distinction is important because human affective ratings reflect lived experience, embodied responses, context, and individual differences that language models do not possess. Nevertheless, prior work suggests that contemporary AI systems can approximate normative affective judgments of images under some conditions (Alrasheed et al., 2025; Romeo & Testolin, 2025; Kim, 2026).

Emotion RDMs derived from AI-generated ratings showed strong correspondence with the human Emotion RDM where the correlation values ranged from 0.70 to 0.79, whereas correspondence among the AI models was higher, with a mean correlation of 0.88. The searchlight fMRI RSA maps derived from the AI Emotion RDMs also converged with the human map in lateral occipitotemporal cortex. At the matched 1,038-voxel level, human–AI overlap was highest for ChatGPT o3 (*J* = .72), followed by Claude 4 Sonnet (*J* = .69) and Gemini 2.5 Pro (*J* = .48). Agreement declined when increasingly selective voxel sets were retained, particularly for Gemini 2.5 Pro. Nevertheless, all observed human–AI overlaps exceeded the permutation null from 100% through 10% retention (one-sided empirical p < .001 at each level).

Together, these findings indicate that a substantial component of the normative valence–arousal structure of the natural scenes considered in this study can be approximated from information captured by current multimodal large language models. This result supports the use of LLM-derived ratings as a complementary resource when human normative ratings are unavailable or when large image sets require rapid characterization. However, the imperfect correspondence with human ratings, the stronger agreement among AI models than between AI and human ratings, and model-specific spatial differences—particularly the additional calcarine cluster for Gemini 2.5 Pro—show that AI-derived ratings should not be regarded as interchangeable with human affective judgments, much less with subjective affective experience.

### Limitations and Future Directions

Several limitations should be noted. First, the study used a relatively small set of 60 standardized affective images. Although the images were selected to span the valence–arousal space, the limited stimulus set may reduce the precision of estimated representational structures and constrain generalizability to other affective categories and more naturalistic materials. Future work should test whether these findings generalize to larger, more diverse image sets and to dynamic stimuli such as videos. Second, the modest participant sample (n = 20) may have limited statistical power to detect weaker or more spatially variable effects and may constrain the generalizability of the observed neural maps. Replication in larger and more diverse samples will be important for assessing the robustness of the reported spatial and temporal patterns. Third, normative valence and arousal ratings do not capture individual variation in affective experience. Future studies could examine whether participant-specific ratings better predict each participant’s neural response patterns. Fourth, the EEG–fMRI fusion analysis establishes a temporal ordering of representational correspondence but does not demonstrate a strictly serial processing sequence or causal flow of information between cortical regions. Finally, LLM-derived ratings depend on model architecture, model version, prompting procedures, and the conditions under which images are presented to the model. They should therefore be viewed as a scalable complement to human normative ratings rather than as substitutes for human affective judgments.

## Notes

https://nda.nih.gov/edit_collection.html?id=2645

### Competing Interest Statement

The authors have declared no competing interest.

### Funding

- NIH Common Fund: MH125615, MH112558

