## Supplement for "Joint Representation of Valence and Arousal in the Human Brain"

### Supplementary Materials

**Definition of Anatomical Regions**

We used HCP-MMP1.0 parcels (Glasser et al., 2016) and the broad regional organization of Horikawa et al. (2020) as a reference to define anatomical regions (Table S1). For example, visual-cortex was broadly divided into MT+, LOC, and remaining VC; the other 13 groups retained their reference memberships. The regions are nonoverlapping, applied separately in each hemisphere, and cover a subset of the atlas that is particularly relevant for the topic examined in this study. A regions was counted as present in a hemisphere if at least one voxel was assigned to one of its parcels.

| **Region** | **Full name** | **Constituent parcels in each hemisphere** |
| --- | --- | --- |
| VC | Visual cortex excluding MT+ and LOC | V1, V2, V3, V3A, V3B, V3CD, V4, V6, V6A, V7, V8, IPS1, FFC, PH, PIT, VMV1, VMV2, VMV3, VVC |
| LOC | Lateral occipital cortex | LO1, LO2, LO3 |
| MT+ | Middle temporal complex | MT, MST, FST, V4t |
| TPJ | Temporo-parietal junction | TPOJ1, TPOJ2, TPOJ3 |
| IPL | Inferior parietal lobule | PFm, PGi, PGs |
| FEF | Frontal Eye Fields | FEF |
| PC | Precuneus | 7m, v23ab, d23ab, 31pv, 31pd, 31a, PCV |
| STS | Superior temporal sulcus | STSda, STSdp, STSva, STSvp |
| TE | Temporal cortex | TE1a, TE1m, TE1p, TE2a |
| MTC | Medial temporal cortex | EC, H |
| Insula | Insular cortex | AAIC, MI, PoI1, PoI2, FOP2, FOP3, Ig, OP2-3 |
| DLPFC | Dorsolateral prefrontal cortex | 9-46d, 46, a9-46v, p9-46v |
| DMPFC | Dorsomedial prefrontal cortex | d32, 9m, 10d |
| VMPFC | Ventromedial prefrontal cortex | 10r, 10v, p32, s32 |
| ACC | Anterior cingulate cortex | a24, a24pr, p24, a32pr, p32pr |
| OFC | Orbitofrontal cortex | OFC, pOFC |

**Table S1.** Anatomical regions and constituent HCP-MMP1.0 parcels in each hemisphere

#### Sensitivity Analysis and Selection of the Searchlight Radius

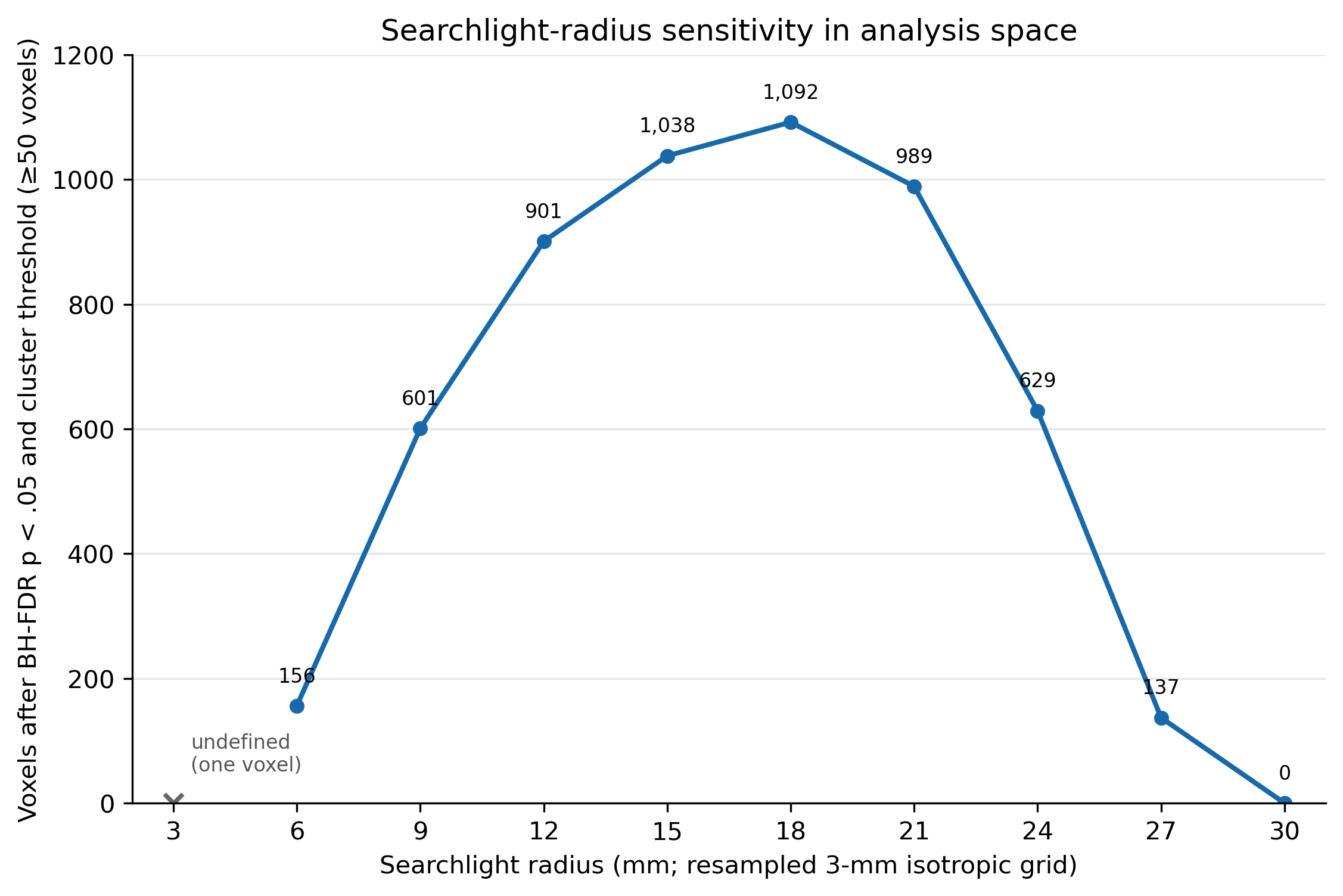

**Figure S1.** Searchlight-radius sensitivity analysis after FDR and cluster-extent thresholding

Functional images were acquired at 3.5-mm isotropic resolution and subsequently resampled to a 3-mm isotropic grid for the searchlight analysis. The primary RSA used a radius of five analysis-space voxels, corresponding to 15 mm and a mean of approximately 484 in-mask voxels per searchlight. This scale was intended to provide sufficient multivoxel information for estimating representational dissimilarity while preserving anatomical specificity. Robustness was assessed by repeating the complete analysis across valid analysis-space radii of 6-30 mm using identical statistical criteria: voxelwise thresholding at FDR-corrected p < .05, restricted to positive effects, followed by a cluster-extent threshold of at least 50 voxels. The final significant extent increased from 156 voxels at 6 mm to 1,038 voxels at the selected 15-mm radius and peaked at 1,092 voxels at 18 mm. Increasing the radius from 15 to 18 mm therefore yielded only 54 additional voxels, or approximately 5%, despite the larger sampling volume. Beyond 18 mm, the significant extent declined to 989 voxels at 21 mm, 629 at 24 mm, 137 at 27 mm, and zero at 30 mm. Because larger searchlights increasingly pool signals across adjacent functional territories and reduce localization precision, maximizing significant-voxel count alone was not considered an appropriate basis for radius selection. The 15-mm radius was therefore retained as a compromise between sensitivity and spatial specificity.

#### Functional Coupling Related to Valence–Arousal Representation and pSTS

We examined functional connectivity among bilateral MT+ (MT, MST, FST, and V4t), LOC (LO1–LO3), TPJ (TPOJ1–TPOJ3), and pSTS (STSdp and STSvp) in 20 participants using beta-series correlations (Rissman et al., 2004). Each ROI included usable voxels within the existing analysis mask. For each participant, hemisphere, and emotion condition (pleasant, neutral, or unpleasant), we averaged the trial-level beta estimates across ROI voxels to obtain 100 values, corresponding to 20 images presented five times. We subtracted each run’s mean and calculated Pearson correlations between ROI beta series. The pSTS series combined STSdp and STSvp, weighted by their usable voxel counts. We transformed the three condition correlations to Fisher-z values and averaged them equally within each participant. Group correlations were obtained by averaging across participants and transforming back to r. Thus, the overall estimate gives equal weight to each condition. Two-sided one-sample t-tests evaluated the 20 participant-level Fisher-z averages against zero (df = 19), with Benjamini–Hochberg FDR correction across the 12 connections in both hemispheres. Ten connections met FDR-adjusted p < .01; right MT+–pSTS and right LOC–pSTS exceeded this threshold.

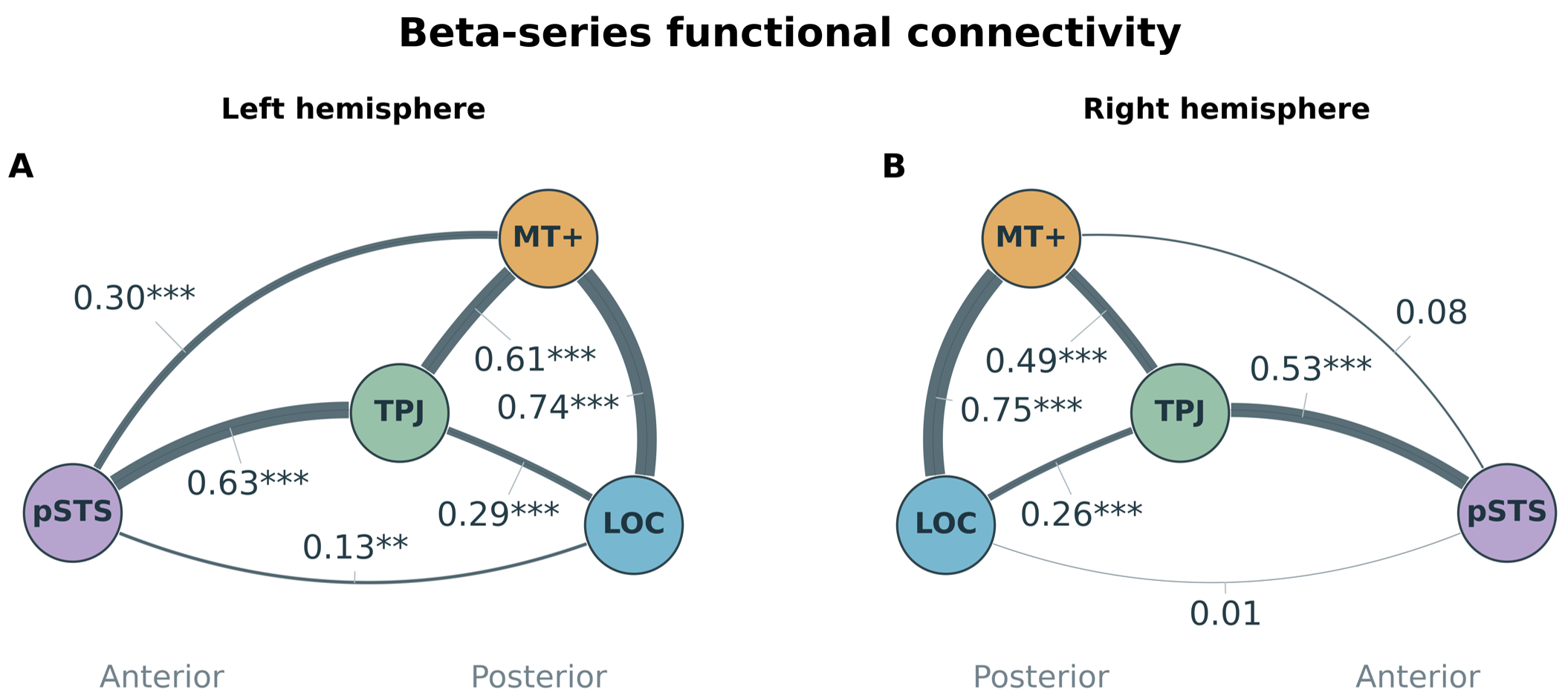

**Figure S2.** Functional coupling related to valence–arousal representation. Correlations among MT+, LOC, TPJ, and pSTS were averaged in Fisher-z space across conditions and 20 participants. Line width reflects r on a fixed 0–1 scale; only positive correlations are shown. Two-sided subject-level tests were FDR-corrected across 12 connections: **p < .01, ***p < .001; unmarked connections did not meet p < .01.

#### LLM Prompts for Valence and Arousal Ratings

The 60 IAPS images were presented to ChatGPT o3, Claude 4 Sonnet, and Gemini 2.5 Pro through their conversational interfaces. Images were uploaded in batches of 10 to accommodate Gemini’s upload limit and maintain a consistent batch size across models. Each model was asked to estimate the ratings that a typical human observer would provide for each image. Valence and arousal were rated on 1-9 scales, with responses reported to two decimal places. Each image was evaluated using three prompts, and the resulting ratings were averaged separately for valence and arousal within each model.

The following is an example prompt:

“For each of the 10 uploaded images, estimate the valence and arousal ratings that a typical human observer would provide. Rate valence from 1 (very unpleasant) to 9 (very pleasant), with 5 indicating neutral valence. Rate arousal from 1 (very calm) to 9 (highly aroused). Report both ratings to two decimal places in a table, identifying each image by its filename or upload order.”
